# Epidermal IL-33 drives inflammation in necroptosis-induced skin inflammation by recruiting TNF-producing immune cells

**DOI:** 10.1101/2023.07.28.550963

**Authors:** Africa Fernandez-Nasarre, Vikas Srivastava, Ffion Bennett, Laurence Michel, Armand Bensussan, Ernest H. Choy, Martine Bagot, Marion C. Bonnet

**Author notes:** These authors contributed equally to this study (Africa Fernandez-Nasarre, Vikas Srivastava). Previous address (Marion C. Bonnet).

## Abstract

Caspase-8 deficiency in the epidermis (caspase-8^EKO^) results in cutaneous inflammation resembling pustular psoriasis, triggered by necroptotic cell death of keratinocytes. Necroptosis is a highly proinflammatory form of programmed necrosis due to the release of intracellular molecules called alarmins, which can act as inflammatory mediators. However, their role in necroptosis-induced skin inflammation remains unexplored. Here, we demonstrate that alarmin IL-33 and its receptor ST2 are essential early mediators of necroptosis-induced skin inflammation. Genetic ablation of *Il-33* or *St2* dramatically delays lesion development and improves survival of caspase-8^EKO^ animals. IL-33 is highly expressed in necroptotic epidermis of caspase-8^EKO^ mice and induces immune cell recruitment in the skin upon keratinocyte necroptosis. Impairment of the IL33-ST2 axis does not affect epidermal necroptosis but reduces the recruitment of TNF-producing infiltrating immune cells and subsequent amplification of cutaneous inflammation. Collectively, our findings highlight a pivotal role for IL-33 and ST2 in necroptosis-induced skin inflammation.

**Teaser:** Inhibition of IL-33/ST2 axis alleviates necroptosis-induced skin inflammation by reducing TNF production in the dermis.

## Introduction

The skin is a complex organ providing a life-sustaining mechanical and chemical barrier to the organism against environmental stresses and pathogens (1). It consists of two main compartments, the dermis and the epidermis. The innermost compartment, the dermis, is mostly constituted of extracellular matrix (collagen, fibronectin) produced by dermal fibroblasts and contains numerous resident immune cells. The outermost compartment, the epidermis, is a stratified epithelium mainly constituted of keratinocytes. Keratinocyte differentiation is a fine-tuned process which is critical to establish the epidermal barrier, which ensures water and temperature homeostasis in mammals. Perturbation of skin homeostasis plays a critical role in chronic inflammatory diseases such as psoriasis or atopic dermatitis (AD) and the crosstalk between dermal and epidermal compartment is pivotal in maintaining the inflammatory state in these diseases (2).

In recent years, keratinocyte cell death has emerged as a critical mechanism in epidermal homeostasis. Cell death imbalance has been shown to trigger skin inflammation in different models. Necroptosis is a programmed form of cell death regulated by Receptor Interacting Protein Kinase 3 (RIPK3) and its substrate Mixed Lineage Kinase domain Like pseudokinase (MLKL) (3). More specifically, necroptotic cell death of keratinocytes has been shown to trigger skin inflammation (4,5). RIPK3/MLKL-dependent necroptosis is responsible for severe skin inflammation in caspase-8 ^EKO^ (6) and FADD ^EKO^ mice (4). While the cutaneous inflammatory phenotype of these mice has been shown to be dependent in part of TNF and TNFR1, the initiating events triggering inflammation in these models remain elusive.

Necroptotic cell death is highly pro-inflammatory due to plasma membrane permeabilisation and subsequent release of pro-inflammatory intracellular molecules, called Damage Associated Molecular Patterns (DAMPs, 7) or alarmins. IL-33 is a cytokine from the IL-1 family. It has initially been identified as a chromatin-associated nuclear factor constitutively expressed in the nuclei of endothelial and epithelial cells (8). It is particularly highly expressed in epidermal keratinocytes. IL-33 has been described as an alarmin or DAMP (9) and acts as a pro-inflammatory mediator upon release from necrotic cells, through binding to its receptor Suppression of Tumorigenicity 2 (ST2). IL-33 has initially been described as a Th2 cytokine (10), due to the high expression of its receptor ST2 on Th2 lymphocytes, mast cells and Innate Lymphoid Cells type 2 (ILC2). IL-33 is also very highly expressed in allergic diseases such as asthma or atopic dermatitis (AD) (11). However, more recently, several studies have suggested a potential role for IL-33 in psoriasis, both in mouse models such as the imiquimod-induced psoriasiform inflammation (12) and in psoriatic patients (13, 14). Here we investigate the role of IL-33/ST2 axis in necroptosis-induced skin inflammation in caspase-8^EKO^ mice by generating caspase-8^EKO^ animals deficient for *Il-33* or its receptor *St2*. Firstly, we show that *Il-33* gene expression is upregulated before lesion development specifically in the epidermis of caspase-8^EKO^ mice. We then demonstrate that *Il-33* or *St2* genetic deficiency strongly delays the development of skin inflammation and significantly increases the survival of caspase-8 ^EKO^ animals. Finally, we show that IL-33 acts as an epidermis-derived inflammatory mediator upon keratinocyte necroptotic cell death and plays together with its receptor ST2 a critical role in recruiting TNF-producing infiltrating immune cells to the dermis.

## Results

### *Il-33* is strongly overexpressed in caspase-8-deficient necroptotic epidermis

*Il-33* has previously been showed to be one of the most highly upregulated genes in the epidermis of caspase-8^EKO^ mice in a microarray analysis (6).

Epidermal keratinocytes are considered as immunocytes due to their cell-autonomous ability to express and release of inflammatory cytokines, such as IL-1β, IL-6 or IL-8. Some of them, also termed alarmins, such as TSLP, IL-1α or IL-33, are specifically released upon necrotic cell death. RIPK3-MLKL-dependent necroptosis has been shown to be responsible for skin inflammation upon keratinocyte-specific deletion of caspase-8 (6) or FADD (4). To investigate the correlation between necroptosis and IL-33 expression, we performed a double immunostaining for Phospho-MLKL (P-MLKL) and IL-33 in skin samples of caspase-8^EKO^ mice and control littermates at postnatal day 1 (P1), P3 and P8. An increase in MLKL phosphorylation could already be detected in the basal and suprabasal layers of caspase-8-deficient epidermis at P1 and the number of P-MLKL positive cells increased at P3 and P8 (Fig. 1).

**Figure 1:**
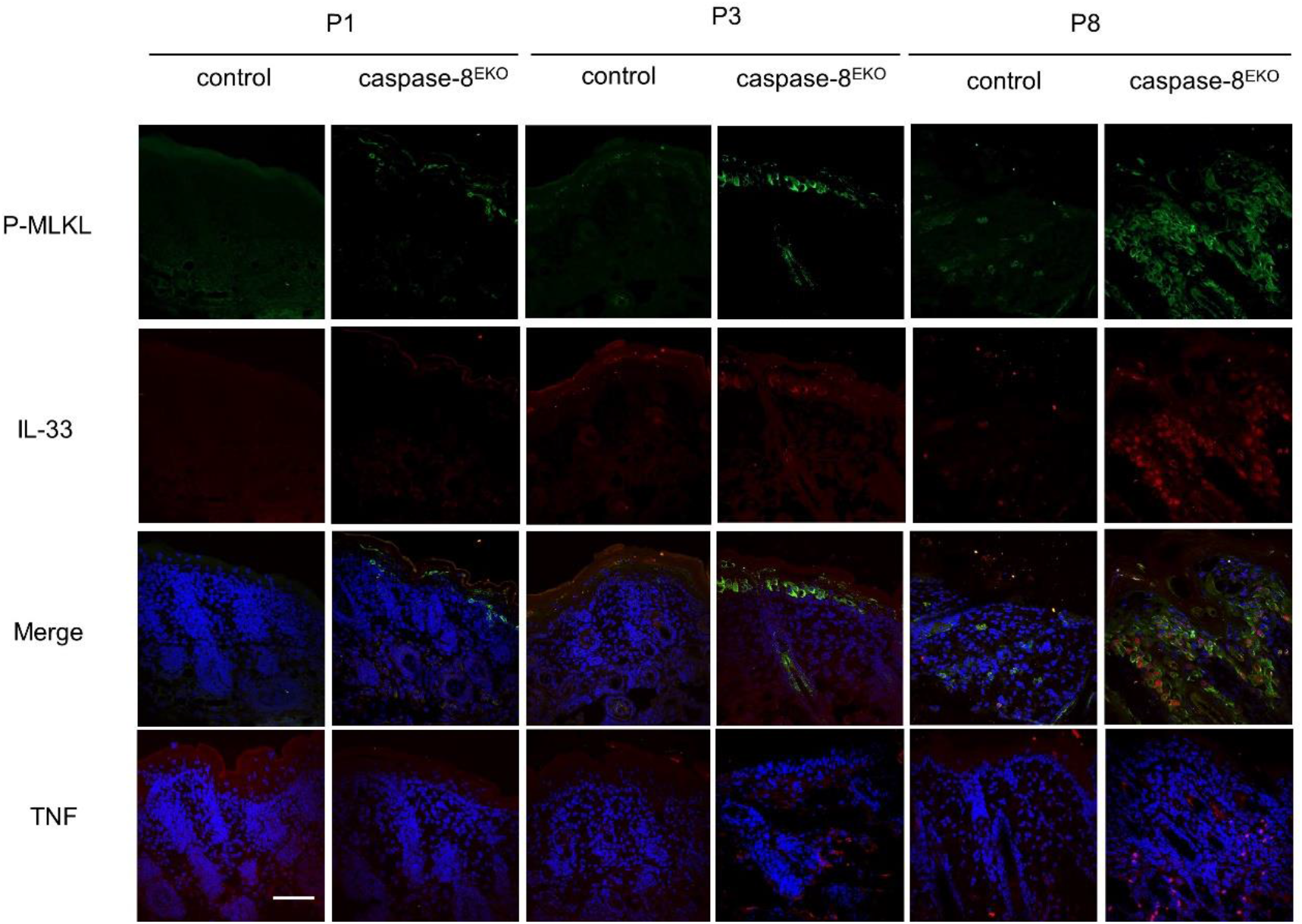
Increased expression of IL-33 from necroptotic keratinocytes in caspase-8^EKO^ mice after birth. Representative images from skin sections of control and caspase-8^EKO^ mice at the indicated ages, immunostained with anti-P-MLKL (green) and anti-IL-33 (red) antibodies and counterstained with DAPI in the merge picture. Lower row: Immunostaining with anti-TNF antibody (red) counterstained with DAPI. P1: control n= 6, caspase-8 ^EKO^ n=6, P3: control n=5, caspase-8 ^EKO^: n=7, P8: control n=3, caspase-8 ^EKO^ n=4. Scale bar=50 μm

At P8, where the majority of the epidermis in caspase-8^EKO^ animals is hyperplastic, we observed that MLKL phosphorylation was clearly associated with lesional areas of epidermal thickening but was absent in non lesional areas of the skin. By comparison, expression of IL-33 could not be found at P1 by immunostaining but was first detected in basal and suprabasal keratinocytes at P3 with the number of IL-33 positive cells increasing with age (Fig. 1). Interestingly, at P8, IL-33 displayed a nuclear localization, while it was mostly found in the cytoplasm at P3. This could indicate a neosynthesis and maybe a cycling of IL-33 expression in hyperproliferative epidermis. IL-33 is expressed specifically in epidermal keratinocytes and was not found in dermal fibroblasts nor in dermal immune cells (Fig. 1). More importantly, co-immunostaining for P-MLKL and IL-33 highlighted that IL-33 expression was strictly restricted to P-MLKL positive keratinocytes at all stages of disease development. This demonstrated that MLKL phosphorylation precedes IL-33 expression.

TNF has been shown to be a major pro-inflammatory mediator in necroptosis-dependent skin inflammation (4,6). Genetic ablation of *Tnf* or *Tnfr1* in FADD^EKO^ or caspase-8^EKO^ mice significantly alleviates skin inflammation in these animals, albeit not fully abrogating it. Hence, we next investigated TNF expression pattern compared to MLKL phosphorylation and IL-33 expression. In agreement with previous reports, TNF expression was strictly restricted to infiltrating immune cells in the dermis and was not detected in the epidermis (Fig.1, bottom row). TNF expression was not detectable in the skin of caspase-8^EKO^ mice at P1. Consistent with previous reports in the literature (4), a few TNF-positive infiltrating cells were first detected at P3 and the number of TNF expressing cells increased in the dermis of caspase-8^EKO^ mice at later timepoints, here shown at P8. This suggests that TNF expression occurs subsequently to keratinocyte necroptosis and IL-33 expression.

Collectively, these results revealed that MLKL is phosphorylated as early as P1 in caspase-8-deficient epidermis and precedes IL-33 expression. Our data also highlighted that IL-33 expression is restricted to P-MLKL-positive keratinocytes. Moreover, our data showed a significant increase in IL-33 expression in the epidermis at P3 at the time when the first TNF-expressing cells can be detected in the dermis of caspase-8^EKO^ mice. Hence, IL-33 is an early marker of skin inflammation upon epidermal necroptosis.

### Differentiation induces *Il-33* expression in mouse keratinocytes and potentiates *Il33* gene induction by cytokines

Our results demonstrate an increase in IL-33 protein expression in caspase-8^EKO^ mice in necroptotic epidermis. To investigate *Il-33* gene expression in caspase-8-deficient epidermis, we performed *Il-33* qRT-PCR analysis on total epidermis from caspase-8^EKO^ mice and control littermates at post-natal day 1 and 3 (P1 and P3). Our results show a significant increase in *Il-33* gene expression of 36-fold at P3 (Fig. 2A). *Il-33* mRNA levels also showed a trend towards increased expression at P1 (3-fold). These results are in agreement with previous reports in the literature (6).

**Figure 2:**
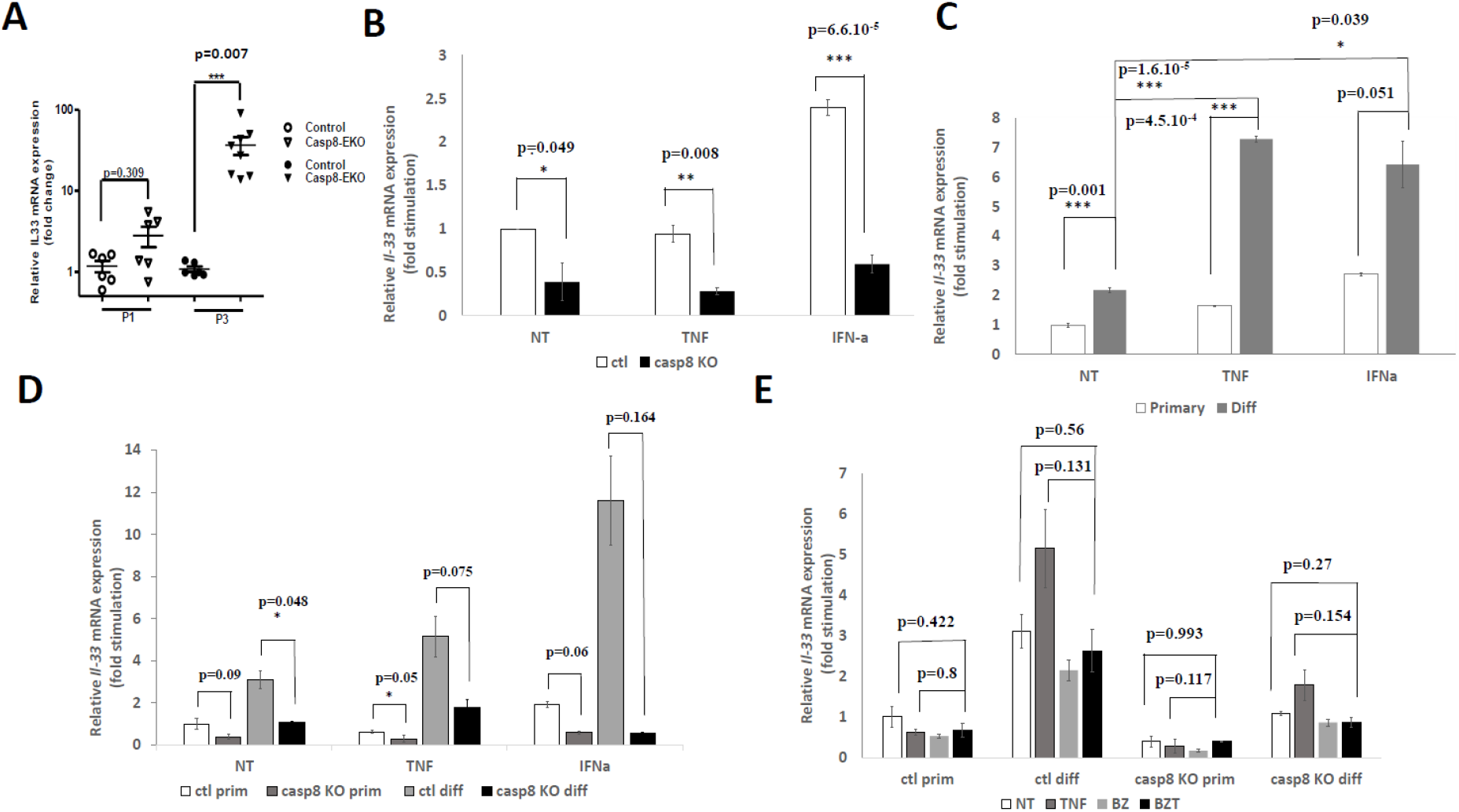
Differentiation but not caspase-8 genetic deficiency induces *Il-33* expression in keratinocytes. A. *Il-33* gene expression in control and caspase-8-deficient epidermis at P1 and P3. Each dot represents individual animals. P1: control: n=6, caspase-8 ^EKO^ n=6, P3: control: n=5, caspase-8 ^EKO^ n=7. B-E: Total RNA was collected 6h after keratinocytes stimulation with mTNF (50 ng/ml) or mIFNα (10^6^ IU). Representative data from min. three independent experiments. B. *Il-33* gene expression in caspase-8 KO primary keratinocytes compared to controls. C. *Il-33* gene expression in control keratinocytes upon keratinocyte differentiation. D. *Il-33* gene expression in caspase-8 KO vs control keratinocytes upon differentiation. E. *Il-33* gene expression upon necroptosis induction in caspase-8 KO vs control keratinocytes. For necroptosis induction, cells were treated for one hour with Smac mimetic BV6 (5 μm) + pan-caspase inhibitor z-VAD (20μm) prior to mTNF stimulation (referred to BZT).

Next, we examined *Il33* gene expression from keratinocytes *in vitro*. Primary keratinocytes were isolated from epidermis of caspase-8 ^EKO^ mice or control littermates at P3 and cultured on collagen-coated plates in low Ca^2+^ culture medium, as previously described (15). Caspase-8-deficient keratinocytes displayed similar growth *in vitro* to control keratinocytes. *Il-33* was constitutively expressed and was detectable at basal level from control epidermal keratinocytes cultured *in vitro*, in agreement with previous reports in the literature (9). Interestingly, caspase-8 deficiency resulted in decreased basal levels on *Il33* gene expression from primary keratinocytes (Fig. 2B). Primary keratinocytes were then differentiated *in vitro* by increasing Ca^2+^ concentration in the culture medium for 20h (1.88 mM CaCl_2_ final). Interestingly, differentiation consistently increased *Il33* gene expression by 2.5-fold in control keratinocytes (Fig. 2C). More importantly, while stimulation with TNF failed to induce *Il33* gene expression in control primary keratinocytes, TNF treatment strongly increased *Il33* mRNA levels in differentiated control keratinocytes by 7-fold (Fig. 2C). Interestingly, only IFNα could stimulate *Il33* gene expression in primary control keratinocytes, which was further enhanced in control differentiated keratinocytes (2.5-fold and 6.5-fold, respectively, Fig. 2C). Hence, our results show that differentiation unlocks *Il33* gene expression in keratinocytes and sensitizes it to modulation by cytokine stimulation.

Caspase-8-deficient keratinocytes showed lower levels of *Il-33* expression in primary keratinocytes. We then assessed *Il-33* gene expression in caspase-8-deficient keratinocytes upon *in vitro* differentiation. Caspase-8-deficient keratinocytes differentiated properly *in vitro*. Caspase-8-deficient differentiated keratinocytes displayed decreased levels of *Il-33* gene expression compared to control differentiated keratinocytes (Fig. 2D), to levels similar to primary keratinocytes. However, differentiation induced *Il-33* gene expression in caspase-8 deficient keratinocytes by 3-fold compared to caspase-8-deficient primary keratinocytes, as observed in control keratinocytes (supplementary Fig. 1). By contrast, *Il-33* gene expression responded to TNF but not to IFNα stimulation in differentiated caspase-8-deficient keratinocytes.

Finally, as IL-33 protein expression in the epidermis colocalizes strictly with P-MLKL, we assessed whether necroptosis induction could directly induce *Il-33* gene expression in control or caspase-8-deficient keratinocytes. To activate necroptosis, we used TNF in the presence of the Smac mimetic BV6 and pan-caspase inhibitor z-VAD-fmk, referred to hereafter as BZT. Our results showed that BZT stimulation does not increase *Il-33* expression in control nor in caspase-8 keratinocytes, irrespective whether they are primary or differentiated (Fig. 2E). Hence, necroptosis-induction in response to TNF does not activate *Il-33* gene expression in keratinocytes. Further investigation will be required to identify the stimulus triggering *Il-33* gene expression in caspase-8-deficient keratinocytes.

### Necroptosis-induced skin inflammation is dependent on IL-33/ST2 signaling in caspase-8^EKO^ mice

To specifically assess the potential role of IL-33 signaling in the development of inflammatory skin lesions in caspase-8^EKO^ mice, we used the genetic mouse model of necroptosis-dependent skin inflammation triggered by keratinocyte-specific genetic ablation of caspase-8 (caspase-8^EKO^ mice, 6) and crossed caspase-8^EKO^ animals with mice deficient for *Il-33* (*Il-33*^-/-^,16) or its receptor *St2* (*St2*^-/-^,17). Caspase-8^EKO^ *Il-33*^-/-^ and caspase-8^EKO^ *St2*^-/-^ mice were born at the expected Mendelian ratio and were macroscopically indistinguishable from their control littermates at birth. *Il-33* or *St2* genetic deficiency resulted in a major delay in the development of cutaneous inflammation in caspase-8^EKO^ mice. Caspase-8^EKO^ *Il-33*^-/-^ and caspase-8^EKO^ *St2*^-/-^ animals displayed only minor macroscopic skin lesions at P9, in contrast with the severely inflamed skin of their caspase-8^EKO^ littermates (Fig. 3A).

**Figure 3:**
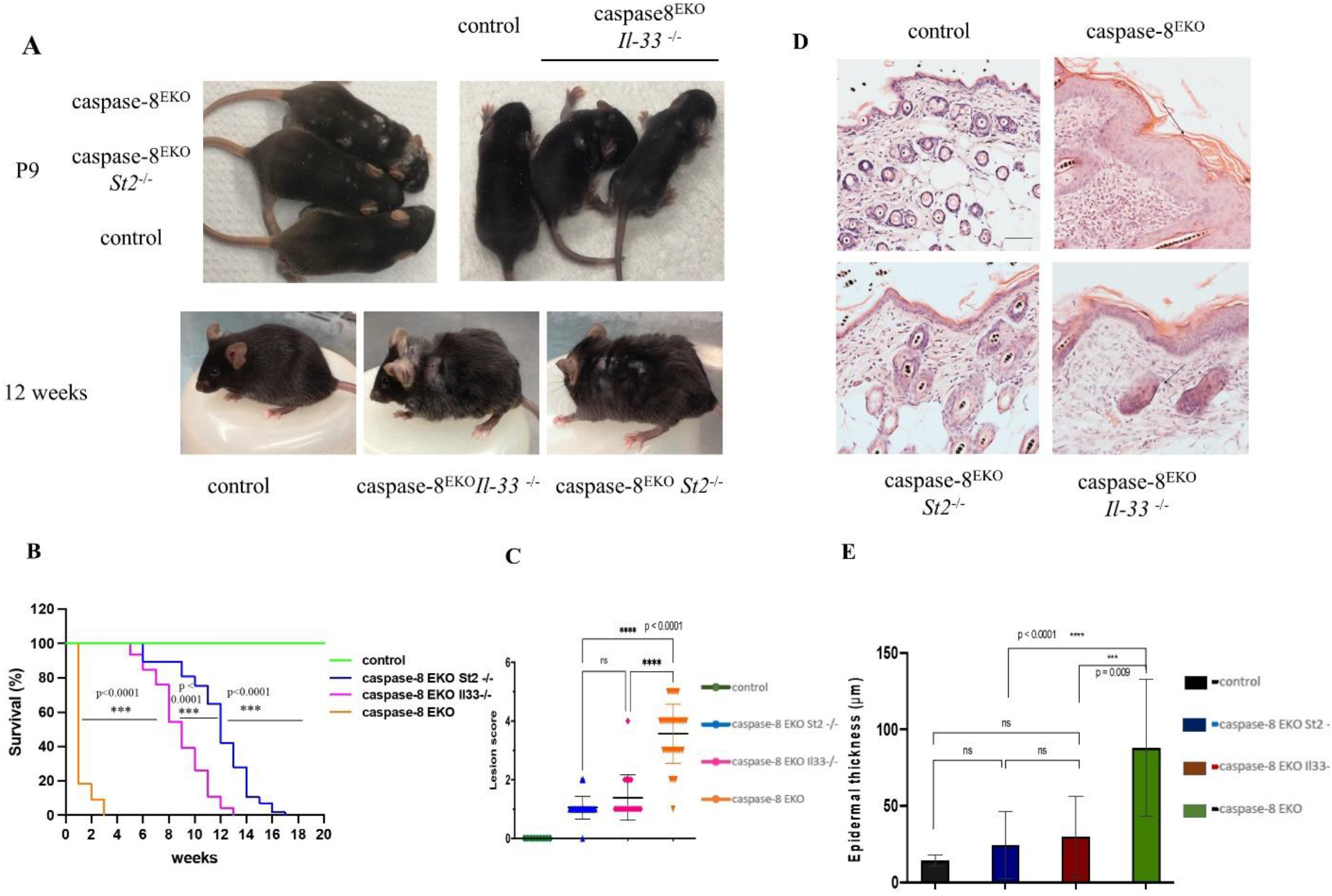
IL-33 or ST2 deficiency alleviate necroptosis-skin inflammation in caspase-8 ^EKO^ mice. A. Representative pictures of mice of the indicated genotypes at P9 and 12 weeks of age. Control: n=27, caspase-8^EKO^ *St2*^*-/-*^: n=57, caspase-8^EKO^ *Il-33*^*-/-*^: n=46, caspase-8^EKO^ =11 B. Kaplan-Meier survival curve of the mice of the indicated genotypes. Control: n=27, caspase-8^EKO^ *St2*^*-/-*^: n=57, caspase-8^EKO^ *Il-33*^*-/-*^ n=46, caspase-8^EKO^ =11 C. Graph depicting macroscopic scoring of skin lesions of the indicated genotypes. Each dot represents an individual mouse. Mean +/-s.e.m. is shown for each group of mice. Statistical significance is determined using ANOVA. Control: n=8, caspase-8^EKO^ *St2*^*-/-*^: n=11, caspase-8^EKO^ *Il-33*^*-/-*^ n=8, caspase-8^EKO^ =9 D. Representative images from of skin sections from control and caspase-8^EKO^ mice at P12 stained with H&E. Control: n=8, caspase-8^EKO^ *St2*^*-/-*^: n=11, caspase-8^EKO^ *Il-33*^*-/-*^ n=8, caspase-8^EKO^ =9. Scale bar=50 μm. E. Graph depicting epidermal thickness measurement of skin sections from mice with the indicated genotype at P12. Each dot represents individual mouse. Mean +/-s.e.m. is shown for each group of mice. Statistical significance is determined using ANOVA. Control: n=8, caspase-8^EKO^ *St2*^*-/-*^: n=11, caspase-8^EKO^ *Il-33*^*-/-*^ n=8, caspase-8^EKO^ =9

Caspase-8^EKO^ *Il-33*^-/-^ and caspase-8^EKO^ *St2*^-/-^ animals showed increased survival up to 13 weeks and 17 weeks respectively compared to 12 days for caspase-8^EKO^ mice (Fig 3A-B and Suppl. Fig. 2). No difference was observed in lesion development nor in survival between males and females.

Macroscopically, inflammation appeared as few isolated patches of inflamed skin at P12, in contrast with widespread skin inflammation in caspase-8^EKO^ animals at this age (Suppl. Fig. 2). The progression of skin inflammation was also strongly delayed compared to caspase-8^EKO^ mice. The lesional score was calculated according to the percentage of body surface affected by the lesions (see Table 1 in Supplementary Material). It was significantly reduced in *Il-33* and *St2*-deficient mice with a similar lesional score of 1 for caspase-8^EKO^ *Il-33*^-/-^ and caspase-8 ^EKO^ *St2*^-/-^ animals at P12 compared to a score of 4.5 for caspase-8^EKO^ mice (Fig 3C).

Histological analysis of sections of back skin of caspase-8^EKO^ mice, caspase-8^EKO^ *Il-33*^-/-^ animals, caspase-8^EKO^ *St2*^-/-^ mice and control littermates at P12 showed a major improvement of epidermal hyperplasia in caspase-8^EKO^ *Il-33*^-/-^ and caspase-8^EKO^ *St2*^-/-^ mice compared to caspase-8^EKO^ animals (Fig 3D). Epidermal thickness measurement at P12 showed a dramatic decrease in epidermal hyperplasia in the skin of caspase-8^EKO^ *Il-33*^-/-^ and caspase-8 ^EKO^ *St2*^-/-^ animals compared to the skin of caspase-8^EKO^ mice, with 90% of skin displaying a normal epidermal thickness, comparable to that of control littermates (Fig. 3E). A slight trend towards increased thickness of the epidermis was observed in caspase-8^EKO^ *Il-33*^-/-^ mice, albeit not significant. However, caspase-8^EKO^ *Il-33*^-/-^ and caspase-8 ^EKO^ *St2*^-/-^ mice ultimately developed generalised skin inflammation after weaning age, with skin lesions affecting 50% of the body surface by 13-17 weeks of age respectively.

Caspase-8^EKO^ *Il-33*^-/-^ and caspase-8^EKO^ *St2*^-/-^ animals displayed very similar phenotypes until 3 weeks of age, with very mild skin inflammation characterized by the presence of very few patches of scaly skin by weaning age. Interestingly, around 3-4 weeks of age, we observed slightly more severe lesions in caspase-8^EKO^ *Il-33*^-/-^ mice and skin inflammation evolved more rapidly compared to caspase-8 ^EKO^ *St2*^-/-^ animals. This resulted in a slight increase in survival of caspase-8^EKO^ *St2*^-/-^ mice compared to caspase-8 ^EKO^ *Il-33*^-/-^ mice. However, both *Il-33* and *St2* genetic ablation significantly improved the skin inflammatory phenotype of caspase-8 ^EKO^ animals, demonstrating a critical role of epidermis-derived IL-33 in necroptosis-dependent cutaneous inflammation.

### Necroptosis is active in lesional epidermis of caspase-8 ^EKO^ *Il33*^-/-^ and caspase-8 ^EKO^ *St2*^-/-^ mice

Skin inflammation in caspase-8^EKO^ and Fadd^EKO^ mice is dependent on RIPK3/MLKL-mediated necroptosis of epidermal keratinocytes (4). To address the impact of IL-33/ST2 signaling on keratinocyte cell death, we then investigated apoptosis and necroptosis, as well as IL-33 expression in the skin of caspase-8^EKO^, caspase-8^EKO^ *Il33*^-/-^ and caspase-8^EKO^ *St2*^-/-^ animals and control littermates by immunofluorescent staining.

As previously described for Fadd^EKO^ mice (4), apoptosis activation marker cleaved caspase-3 was not detected in dying keratinocytes in the epidermis of caspase-8^EKO^, caspase-8^EKO^ *Il33*^-/-^ and caspase-8^EKO^ *St2*^-/-^ mice, showing that epidermal keratinocytes do not undergo apoptotic death in these animals (supplementary Fig.3A).

Next, we assessed the activation of the necroptotic pathway through immunostaining of the necroptotic central regulatory molecule, protein kinase RIPK3 (Supplementary Fig. 3B). Firstly, we observed a strong increase in RIPK3 expression in the epidermis of caspase-8^EKO^ littermates at P12. RIPK3 expression was particularly prominent in keratinocytes in hyperplastic lesional epidermis, but RIPK3 was not detected in non lesional epidermis. Interestingly, RIPK3 expression was also strongly upregulated in the epidermis of caspase-8^EKO^ *Il33*^-/-^ and caspase-8 ^EKO^ *St2*^-/-^ animals. We therefore hypothesized that necroptosis might still be active in the absence of IL-33/ST2 signaling.

We then investigated necroptosis activation through immunostaining of RIPK3 substrate, phosphorylated MLKL. We also assessed IL-33 expression in the epidermis of caspase-8^EKO^, caspase-8 ^EKO^ *Il33*^-/-^ and caspase-8 ^EKO^ *St2*^-/-^ animals at P12 (Fig. 4).

**Figure 4:**
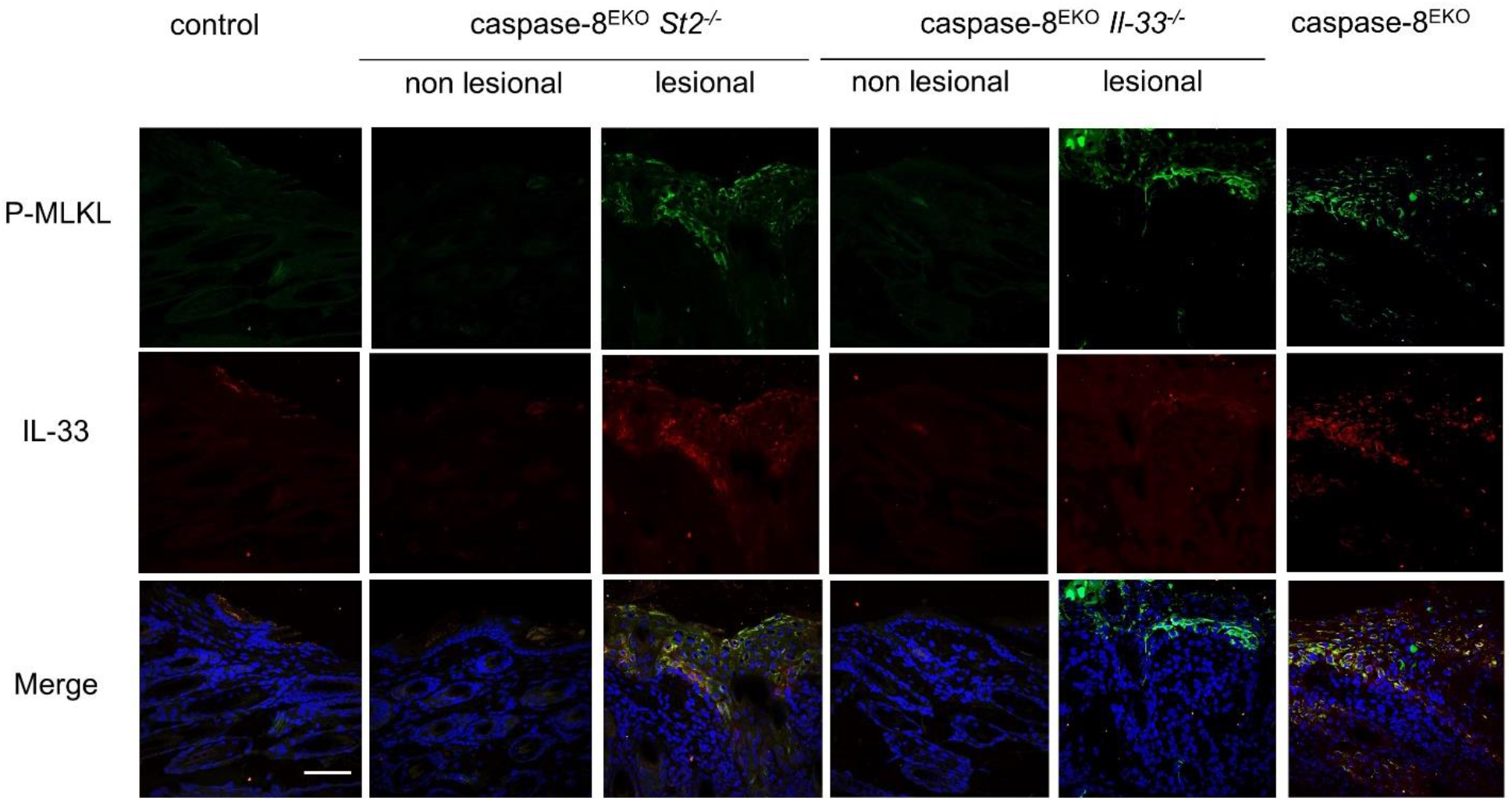
IL-33 and ST2 deficiency do not prevent necroptosis in caspase-8-deficient epidermis. Representative images from skin sections of mice of the indicated genotypes, immunostained with anti-P-MLKL (green) and anti-IL-33 (red) antibodies and counterstained with DAPI (blue) in the merge picture. Control: n=6, caspase-8^EKO^ *St2*^*-/-*^: n=6, caspase-8^EKO^ *Il-33*^*-/-*^ n=5, caspase-8^EKO^ =5. Scale bar=50 μm.

As observed at P8, caspase-8-deficient epidermis displayed a strong immunostaining for P-MLKL at P12 in hyperplastic lesions. IL-33 was also strongly expressed in lesional epidermis and colocalised strictly with P-MLKL. Of note, IL-33 was not detected from immune cell infiltrates in the dermis. Interestingly, the skin of caspase-8^EKO^ *Il33*^-/-^ and caspase-8^EKO^ *St2*^-/-^ animals still showed increased levels of phosphorylation of MLKL in lesional areas of the epidermis but P-MLKL could not be detected in non lesional areas. As expected, IL-33 immunostaining was totally absent in caspase-8^EKO^ *Il33*^-/-^ animals. However, IL-33 was still expressed in the epidermis of caspase-8^EKO^ *St2*^-/-^ mice. As observed in caspase-8^EKO^ animals (Fig. 1 and 4), IL-33 expression was restricted to P-MLKL positive cells in the epidermis of caspase-8^EKO^ *St2*^-/-^ animals.

Our data demonstrated that abrogation of IL-33/ST2 signaling does not affect keratinocyte necroptotic cell death. Increased IL-33 expression in the epidermis of caspase-8^EKO^ mice was also not inhibited by *St2* genetic ablation. Taken together, IL-33/ST2 axis mediates skin inflammation downstream of keratinocyte necroptosis.

### Genetic ablation of *Il33 or St2* restores epidermal differentiation in caspase-8^EKO^ mice

Our results have shown that genetic ablation of *Il33* or *St2* significantly improves epidermal hyperplasia in caspase-8^EKO^ mice (Fig. 3). However, necroptosis marker P-MLKL could still be detected in the epidermis of caspase-8^EKO^ *Il33*^-/-^ and caspase-8^EKO^ *St2*^-/-^ mice. Given the persistence of necroptosis in caspase-8 deficient epidermis, we next assessed whether necroptosis had an impact on epidermal differentiation in the skin of caspase-8^EKO^ *Il33*^-/-^ and caspase-8^EKO^ *St2*^-/-^ animals. For this purpose, we analysed the expression of epidermal differentiation markers in skin sections of caspase-8^EKO^ *Il33*^*-*/-^ and caspase-8^EKO^ *St2*^-/-^ mice by comparison with the skin of caspase-8^EKO^ and control littermates at P12 (Fig. 5).

**Figure 5:**
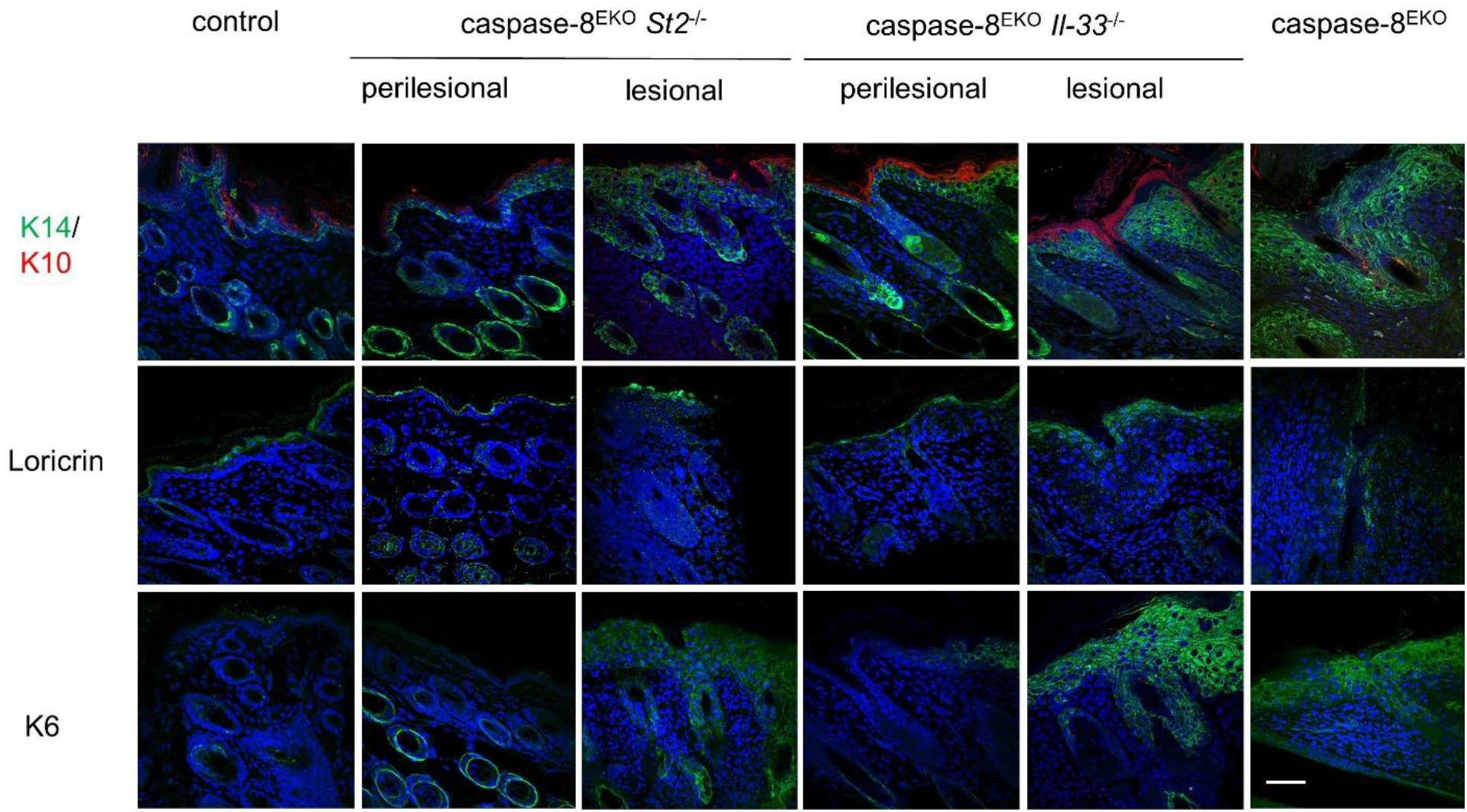
IL-33 and ST2 deficiency restores normal epidermal differentiation on most of the skin surface. Representative images from skin sections of the mice of the indicated genotypes, immunostained with anti-K14 (green) and anti-K10 (red), anti-loricrin and anti-K6 antibodies and counterstained with DAPI. Control: n=4, caspase-8^EKO^ *St2*^*-/-*^: n=4, caspase-8^EKO^ *Il-33*^*-/-*^ n=5, caspase-8^EKO^ n=3. Scale bar=50 μm.

Immunostaining with antibodies specific for basal layer marker keratin 14 (K14) and suprabasal marker keratin 10 (K10) revealed a similar pattern of expression of K14 and K10 in the non lesional skin of caspase-8^EKO^ *Il33*^*-*/-^ and caspase-8^EKO^ *St2*^-/-^ animals compared to the differentiation pattern of control skin (Figure 5). K14 expression was progressively increased in perilesional areas of caspase-8^EKO^ *Il33*^*-*/-^ and caspase-8^EKO^ *St2*^-/-^ mice skin while K10 expression was maintained. However focal areas of epidermal hyperplasia in the skin of caspase-8^EKO^ *Il33*^*-*/-^ and caspase-8^EKO^ *St2*^-/-^ mice were characterized by ubiquitous expression of K14 throughout all layers of hyperplastic epidermis and a decreased expression of K10.

We next assessed the expression of the granular layer marker loricrin in the epidermis of caspase-8^EKO^ *Il33*^*-*/-^ and caspase-8^EKO^ *St2*^-/-^ mice. Similarly, loricrin expression was maintained in the non lesional epidermis of caspase-8^EKO^ *Il33*^*-*/-^ and caspase-8^EKO^ *St2*^-/-^ mice. Loricrin expression was also maintained in perilesional and lesional skin in caspase-8^EKO^ *Il33*^*-*/-^ and caspase-8^EKO^ *St2*^-/-^ animals, by contrast with the loss of expression of loricrin in the epidermis of caspase-8^EKO^ animals. In lesional areas of caspase-8^EKO^ *Il33*^*-*/-^ and caspase-8^EKO^ *St2*^-/-^ epidermis, the loricrin positive granular layer appeared thicker than in non lesional areas, displaying two to three cell layers compared to a single cell layer in non lesional areas. Finally, we analysed the expression of Keratin 6 (K6), an epidermal stem cell marker also increased in hyperproliferative epidermis. In non lesional skin of caspase-8^EKO^ *Il33*^*-*/-^ and caspase-8^EKO^ *St2*^-/-^ mice, K6 expression was restricted to hair follicle as observed in control skin. Areas of focal epidermal hyperplasia correlated with increased expression of K6 in all layers of the interfollicular epidermis, as observed in caspase-8 deficient epidermis.

Taken together, these results are consistent with the macroscopic observation and histological analysis showing only focal areas of skin inflammation in caspase-8^EKO^ *Il33*^*-*/-^ and caspase-8^EKO^ *St2*^-/-^ animals at P12 (Fig. 3A and D). Our observations show that *Il-33*/*St2* deficiency in caspase-8^EKO^ mice restores a normal differentiation pattern in 90% of the epidermis, similar to the pattern observed in control littermates, but that focal areas of epidermal hyperplasia in the skin of caspase-8^EKO^ *Il33*^*-*/-^ and caspase-8^EKO^ *St2*^-/-^ animals display characteristic hallmarks of hyperplastic epidermis as observed in the skin of caspase-8^EKO^ animals. Finally, these results demonstrate that altered epidermal differentiation is not caused by increased keratinocyte necroptosis itself but by the subsequent proinflammatory cascade triggered by the release of IL-33 and activation of ST2 signaling.

### Inhibition of IL-33 signaling impairs immune cell recruitment and TNF production in necroptotic epidermal lesions

Cutaneous inflammation and epidermal hyperplasia in caspase-8^EKO^ mice is associated with the presence of immune cell infiltrates in the dermis, comprising T cells, macrophages and granulocytes (6,4). To address the role of IL-33/ST2 signaling in immune cell recruitment, we investigated the immune cell population present in the dermis in caspase-8^EKO^ *Il33*^*-*/-^ and caspase-8^EKO^ *St2*^-/-^ mice. Immunostaining of skin sections was performed using specific markers of T cells (CD3), macrophages (F4/80) and granulocytes (Gr-1). As described previously, increased infiltration of T cells, macrophages and granulocytes was observed in the dermis of caspase-8^EKO^ mice at P12 (Fig.6).

**Figure 6:**
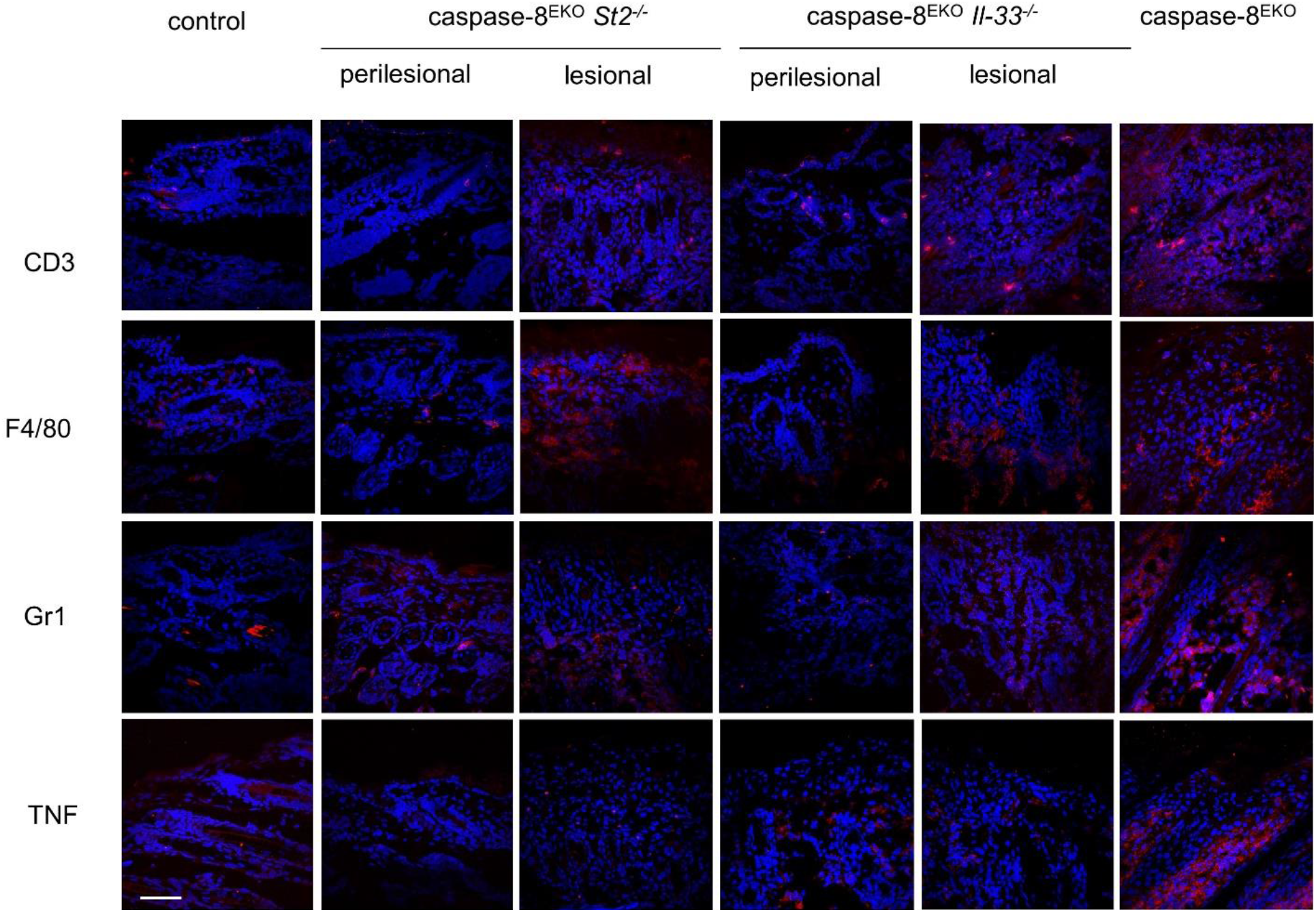
*Il-33* and *St2* deficiency limit the recruitment of TNF-producing infiltrating immune cells in the skin upon epidermal necroptosis. Representative images of skin sections from mice of the indicated genotypes, immunostained with anti-CD3, anti-F4/80, anti-Gr-1 or anti-TNF antibodies and counterstained with DAPI. Control: n=6, caspase-8^EKO^ *St2*^*-/-*^:n=6, caspase-8^EKO^ *Il-33*^*-/-*^ n=5, caspase-8^EKO^ =4. Scale bar=50 μm.

Interestingly, we observed similar numbers of CD3-positive T cells and F4/80-positive macrophages, in the lesional skin of caspase-8^EKO^ *Il33*^-/-^ and caspase-8^EKO^ *St2*^-/-^ mice, suggesting that recruitment of T cells and macrophages is unaffected upon genetic ablation of *Il-33* or *St2* compared to caspase-8^EKO^ mice. In contrast, the number of infiltrating Gr-1 positive granulocytes was significantly reduced in the skin of *Il-33* or *St2*-deficient animals compared to the skin of caspase-8^EKO^ mice. Hence, IL-33/ST2 axis appears to be specifically involved in granulocyte recruitment at the site of the lesion but does not affect T cells nor macrophages infiltration upon keratinocyte necroptotic cell death.

Skin inflammation in caspase-8^EKO^ mice has been shown to be strongly dependent upon TNF/TNFR1 signaling (4,6). To assess the impact of IL-33/ST2 signaling on cutaneous TNF expression, skin cryosections from caspase-8 ^EKO^, caspase-8 ^EKO^ *Il33*^-/-^ and caspase-8^EKO^ *St2*^-/-^ mice and control littermates were stained with an antibody specific for murine TNF. In caspase-8^EKO^ mice skin, TNF expression was found exclusively in the dermis, with a distribution pattern similar to the distribution of immune cell infiltrates. TNF immunostaining revealed a significant decrease in TNF expression in the skin of caspase-8^EKO^ *Il33*^*-*/-^ and caspase-8 ^EKO^ *St2*^-/-^ mice compared to caspase-8^EKO^ mice. Our results demonstrate that IL-33/ST2 axis controls TNF abundancy in the dermis upon epidermal necroptosis. Taken together, our data show that IL-33/ST2 deficiency had little impact on T cells and macrophages numbers in the dermis but inhibited specifically granulocytes recruitment and TNF production in caspase-8^EKO^ necroptosis-dependent skin inflammation.

## Discussion

Necroptosis has gained increasing scientific interest as an initiating mechanism of inflammatory diseases since the first demonstration of the role of keratinocyte necroptosis as a trigger for cutaneous inflammation in FADD^EKO^ mice (4). Epidermis appears particularly susceptible to necroptosis, as evidenced by several models of necroptosis-driven skin inflammation upon keratinocyte-specific deletion of RIPK1, NF-κB subunits or more recently OTULIN (5, 32-35). Yet, the inflammatory mediators responsible for necroptosis-dependent skin inflammation remain to be identified.

Here we provide evidence that the IL-33/ST2 axis is a critical initiator of necroptosis-induced skin inflammation in caspase-8^EKO^ mice. *Il-33* gene expression has been shown to be upregulated in the epidermis of caspase-8^EKO^ animals shortly after birth (6). IL-33 expression was found exclusively in the caspase-8-deficient epidermis, but not in the dermis, and was restricted to P-MLKL-positive necroptotic keratinocytes (Fig.1). *Il-33* gene expression was found to be upregulated in caspase-8-deficient epidermis already at P1 (Fig.2A). However, in agreement with previous studies (6), no increase in *Il-33* gene expression was found in primary caspase-8 KO keratinocytes in culture compared to control keratinocytes (Fig. 2B). Interestingly, we show here that *Il-33* gene expression is increased by three-fold upon keratinocyte differentiation *in vitro* in both control and caspase-8 KO keratinocytes (Fig. 2C-D and Suppl. Fig. 1). This confirms that *Il-33* upregulation is not an intrinsic property of caspase-8 KO keratinocytes. More interestingly, while *Il-33* expression was not altered by cytokine stimulation in control or caspase-8 KO primary keratinocytes, *Il-33* expression was significantly increased by TNF or IFNα in differentiated keratinocytes. However, again, *Il-33* expression was still not stronger in caspase-8-deficient differentiated keratinocytes upon cytokine stimulation. Despite IL-33 protein expression being restricted to P-MLKL-positive necroptotic keratinocytes, necroptosis induction by TZB did not enhance but rather decreased *Il-33* expression in primary and differentiated keratinocytes *in vitro*, irrespective of their genotype (Fig. 2E). Hence, *Il-33* gene expression was not directly upregulated by necroptosis induction in control or caspase-8-deficient keratinocytes *in vitro*. Further investigation will be needed to identify factors leading to increased *Il-33* expression in caspase-8-deficient epidermis.

In order to assess the role of IL-33 and its receptor ST2 in necroptosis-dependent cutaneous inflammation *in vivo*, we have generated caspase-8^EKO^ mice with genetic deficiency for *Il-33* or its receptor *St2*. Genetic ablation of either *Il-33* or *St2* significantly alleviated skin inflammation in caspase-8^EKO^ mice, resulting in a major improvement of the survival of these animals (Fig. 3A-B). Normal epidermal thickness was restored on 90% of the body surface (Fig. 3D-E) and. However, caspase-8 ^EKO^ *Il-33*^-/-^ and caspase-8^EKO^ *St2* ^-/-^ mice do present focal areas of cutaneous inflammation. Interestingly, the development of the lesions is much slower in caspase-8^EKO^ *Il-33*^-/-^ and caspase-8^EKO^ *St2*^-/-^ mice than in caspase-8 ^EKO^ mice and the progression of the disease is delayed. However, cutaneous inflammation does not resolve, and animals reached the experimental endpoint of 50% body surface affected by skin lesions at 13 weeks of age for caspase-8^EKO^ *Il-33*^-/-^ mice and 17 weeks of age for caspase-8^EKO^ *St2*^-/-^ animals.

We also investigated cell death markers in caspase-8 ^EKO^ *Il-33*^-/-^and caspase-8 ^EKO^ *St2*^-/-^ mice. Interestingly, *Il33* or *St2* deficiency does not prevent keratinocyte necroptosis, which was found to be still active in focal areas of skin lesions in caspase-8 ^EKO^ *Il-33*^-/-^ and caspase-8 ^EKO^ *St2*^-/-^ animals (Fig.4). Additionally, IL-33 expression was also increased in caspase-8 EKO *St2*^-/-^ mice in necroptotic keratinocytes at the site of lesions. This suggests that the pro-inflammatory IL-33/ST2 cascade acts downstream of keratinocyte necroptosis in the course of cutaneous lesion development.

We further investigated immune cell infiltration in caspase-8^EKO^ *Il-33*^-/-^ and caspase-8^EKO^ *St2*^-/-^ mice. No significant difference was observed in the recruitment of CD3^+^ T cells. This is in agreement with previous data showing that lymphocyte depletion through *Rag1* genetic ablation does not rescue necroptosis-induced skin inflammation in Fadd^EKO^ animals (4). Similarly, F4-80^+^ macrophages abundancy was also not altered in skin lesions upon genetic ablation of *Il-33* or *St2* in caspase-8 ^EKO^ mice. This is consistent with the observation that macrophage depletion using clodronate liposomes did not alleviate skin inflammation in caspase-8 ^EKO^ animals (6). In contrast to T cells and macrophages, a significant decrease was observed in Gr-1^+^ granulocytes infiltrate at the site of lesions in the dermis of caspase-8^EKO^ *Il-33*^-/-^ and caspase-8^EKO^ *St2*^-/-^ mice compared to caspase-8^EKO^ mice. Granulocytes constitute a very heterogeneous group of myeloid cells, comprising neutrophils, basophils, eosinophils and mast cells. All subtypes of granulocytes can be found in the skin and can contribute to skin inflammation through the production of pro-inflammatory cytokines, including TNF. Interestingly, IL-33 has been shown to stimulate neutrophil recruitment through induction of TNF production in mast cells (20).

Previous studies have shown that TNF and TNFR1 play an important role in the development of skin inflammation in FADD^EKO^ and caspase-8^EKO^ mice (4,6). However, we have shown that TNF/TNFR1 was not the initial trigger of necroptosis in caspase-8^EKO^ or FADD^EKO^, as genetic ablation of *Tnfr1* did not fully rescue cutaneous inflammation in a similar manner as *Ripk3* deletion does. In line with this observation, TNF expression was found at the site of immune cell infiltration in the dermis, while IL-33 expression colocalized specifically with necroptotic keratinocytes in the epidermis. However, lesion development is only moderately delayed in caspase-8^EKO^ or Fadd^EKO^ mice deficient for *Tnf* or *Tnfr1*. By comparison, genetic ablation of *Il33* or *St2* improve more significantly cutaneous inflammation and animal survival in caspase-8^EKO^ mice than genetic ablation of *Tnf* or *Tnfr1*, with survival rates of 13-17 weeks for caspase-8 ^EKO^ *Il-33*^-/-^ and caspase-8 ^EKO^ *St2*^-/-^ mice and 5-10 weeks for caspase-8 ^EKO^ *Tnf*^-/-^ animals, as well as Fadd ^EKO^ *Tnf*^-/-^ and Fadd ^EKO^ *Tnfr1*^-/-^ mice (4, 6). Moreover, IL-33 expression could be earlier and at the site of necroptotic lesions in the skin compared to TNF expression. Collectively, these results demonstrate that IL-33 expression precedes immune cell infiltration and TNF production. This is strongly supported by the significant decrease in TNF expression in the skin of caspase-8 ^EKO^ *Il-33*^-/-^ and caspase-8 ^EKO^ *St2*^-/-^ mice. The improved survival of caspase-8^EKO^ *Il-33*^-/-^ and caspase-8^EKO^ *St2*^-/-^ animals compared to caspase-8^EKO^ *Tnf*^-/-^ or Fadd^EKO^ *Tnf*^-/-^ and Fadd^EKO^ *Tnfr1*^-/-^ would also suggest that other cytokines than TNF participate to the pro-inflammatory effect of IL-33 and ST2 in the skin upon keratinocyte necroptosis.

We show here that IL-33/ST2 axis is essential to the recruitment of infiltrating immune cells and TNF production. It remains unclear whether IL-33 acts through ST2 as a chemoattractant for TNF-producing cells or if it directly induces TNF production from ST2 positive cells or both. IL-33 has originally been implicated in Th2 response in allergic diseases such as asthma in the lung or Atopic Dermatitis (AD) (11) in the skin. Skin inflammation in our model is strongly dependent on TNF/TNFR1 and macroscopically and histologically resembles psoriasis. However, IL-33/ST2 axis appears to be essential for TNF production in the dermis in our model, which could suggest a role for IL-33 and ST2 in TNF-dependent skin inflammatory conditions, such as psoriasis. Interestingly, increased expression of IL-33 has also recently been described in psoriasis. Serum levels of IL-33 have been shown to be elevated in psoriatic patients (14) and polymorphisms of the IL33 gene have been associated to increased susceptibility to develop psoriatic arthritis (13). ST2 deficiency has been shown to inhibit skin inflammation in the psoriasis model of imiquimod-induced skin inflammation (12). Moreover, IL-33 has been shown to contribute to skin inflammation in mice in a phorbol-ester model of skin inflammation (18). In this model, skin inflammation is partially mediated by mast cells, but IL-33 also triggers the recruitment of neutrophils. Interestingly, another study reported a role for IL-33 in inducing TNF and IL-6 expression from mast cells through a MAPK and PI3K-dependent mechanism resulting in neutrophil recruitment (19). Our data support a potential role of IL-33 and ST2 in the recruitment of granulocytes and induction of TNF in necroptosis-induced skin inflammation.

While the role of IL-33 as an alarmin has been broadly reported, the potential role of necroptosis in *Il-33* gene induction is still unclear. The mechanisms by which necroptosis of a small number of epidermal keratinocytes can trigger the development of severe skin inflammation remained to date elusive. It remains unclear whether inflammatory gene induction could be dependent on necroptotic machinery and necroptotic cell death completion or if it is only concomitant to necroptosis. For example, it has been shown that RIPK3-dependent pro-inflammatory cytokine production induced during necroptosis can persist after membrane permeabilization in ER-containing necroptotic corpses (27). A role for RIPK3 in cytokine induction independent of its pro-necroptotic function has been suggested. Interestingly, another study reported a necroptosis-induced MLKL-dependent increase in cytokine transcription in human and murine cells (28). Another report described that IL-33 protein release was dependent on MLKL function in the airway epithelium but did not mention *Il-33* gene induction in this model (29). Here we show that IL-33 protein expression is restricted to P-MLKL-positive keratinocytes in the epidermis, and that *Il-33* gene expression is increased in caspase-8-deficient epidermis. However, we observed that BZT-induced necroptotic cell death failed to stimulate *Il-33* gene expression in wt or caspase-8-deficient keratinocytes (Fig. 2E). Hence further in-depth investigation is required to identify the regulators of *Il-33* gene expression in caspase-8 KO epidermal keratinocytes.

In summary, our study has identified epidermis-derived IL-33 and its receptor ST2 as central mediators of inflammation in necroptosis-induced skin inflammation, where they contribute to the recruitment of TNF-producing infiltrating immune cells and subsequent amplification of necroptosis-induced inflammation,

## Materials and Methods

### Experimental study design

This study aims at unraveling the role of IL-33/ST2 signaling axis in necroptosis-dependent skin inflammation. Necroptosis activation was assessed by immunostaining of Phospho-MLKL (P-MLKL) in correlation with IL-33 and TNF expression in back skin frozen sections from control and caspase-8^EKO^ mice at post-natal day 1, 3 and 8 (P1, P3 and P8). To investigate the *in vivo* role of IL-33 and its receptor ST2 in necroptosis-induced skin inflammation, we generated caspase-8^EKO^ mice deficient for either *Il-33* or *St2* and studied the impact of *Il-33* or *St2* gene ablation on the development of skin inflammation. Mice were bred in the animal facility at Cardiff University in Specific Pathogen Free conditions. All animal experiments were conducted according to the animal experimentation regulation of the UK Home Office as well as European regulations and were approved by the Home Office and the local Ethics Review Committee at Cardiff University. Caspase-8 flox/flox mice were a kind gift from Pr. Stephen Hedrick (University of California San Diego, U.S.A.). *St2*^-/-^ mice were a kind gift of Pr. Daniel Pinschewer (University of Basel, Switzerland, 17). *Il-33*^-/-^ animals were a kind gift from Dr. Andrew McKenzie (MRC Laboratory for Molecular Biology, Cambridge, U.K., 16) and were provided by Dr. James MacLaren (Cardiff University). K14-Cre transgenic mice were purchased from Jackson laboratories. Macroscopic characterization of the skin inflammatory phenotype was performed by monitoring survival and lesion scoring at different time points. For inflammation monitoring and survival studies, the experimental endpoint was reached when skin lesions affected 50% of the total body surface. For assessment of cell death, epidermal differentiation, and inflammatory markers in caspase-8^EKO^ *Il-33* ^-/-^ and caspase-8^EKO^ *St2*^-/-^ animals, skin samples were taken at P12, which was the experimental endpoint for caspase-8^EKO^ animals. Pathological and clinical parameters of the dorsal skin were assessed by histological analysis, immunofluorescent staining, and confocal imaging. For keratinocyte culture and mRNA isolation, total epidermis was isolated at P1 and P3. The induction of *Il-33* gene expression was characterized by qRT-PCR in total necroptotic epidermis and in cultured mouse keratinocytes. Sample sizes for animal studies were estimated according to similar previous studies. Experiments with statistical analysis were performed in triplicates in at least three independent experiments.

### Isolation of epidermal sheets

Animals were sacrificed at P3, and skin washed briefly in 100% ethanol then in sterile PBS. Total skin was isolated and incubated 30 min at 37 °C in 2.5% trypsin for epidermis RNA isolation or overnight at 4 °C in 0.25 % Trypsin for keratinocyte culture. Following trypsin treatment, the dermis was removed from epidermis. For total epidermis RNA isolation, epidermal sheets were snap frozen and stored at -80 ° until RNA extraction.

### Mouse keratinocyte culture

Mouse primary keratinocytes were isolated from epidermis of caspase-8^EKO^ mice and control littermates collected at P3. Keratinocytes were cultured in an incubator at 5% CO2 at 35°C in 6-well plates (Falcon) coated with PureCol bovine collagen I solution (Cell Systems) in low calcium homemade culture medium containing recombinant mEGF (Peprotech) + Chelex treated FCS (see Supplementary material and 15). Chelex resin was purchased from Biorad. Medium was changed 24h after plating to removed unattached keratinocytes and primary keratinocytes were left to grow until 80% confluence. For *in vitro* differentiation, cells were incubated for 20h prior to stimulation in homemade culture medium supplemented with high CaCl_2_ (1.88 mM final).

### Cytokine stimulation

Primary and/or differentiated keratinocytes were treated for 6h with 50 ng/ml mTNF (Peprotech) or 10^6^ IU mIFNa (PBL Assay Science). For induction of necroptosis, cells were pre-treated for one hour with the Smac mimetic BV6 (5 μM, Selleckchem) + pan-caspase inhibitor zVAD-fmk (20 μM, Selleckchem) prior to mTNF stimulation.

### RNA extraction and qRT-PCR

Total RNA was extracted from total mouse epidermis using TRIzol reagent (Life Technologies) and RNeasy Mini RNA isolation kit (Qiagen) according to manufacturer’s instructions. For *in vitro* cultured keratinocytes, RNA was extracted from two wells of 6-well plates using TRIzol and RNeasy Mini RNA isolation kit, as described above. cDNA synthesis was performed using 1μg of RNA from total epidermis or 500 ng of RNA from keratinocytes with SuperScript™ III First-Strand Synthesis System (Invitrogen). Real-time quantitative RT-PCR (qRT-PCR) was carried out in triplicate in a 96-well plate using PowerUp™ SYBR™ Green Master Mix (Applied Biosystems™) in a QuantStudio ViiA7 Flex real-time PCR system (Applied Biosystems™). qRT-PCR for *mIl-33* and *mGapdh* primers are described below. Data were quantified using 2^ΔΔCT^ method.

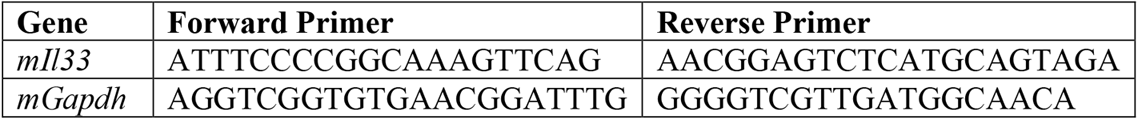

### Tissue lysis and genotyping protocols

For animal genotyping, ear punch biopsies were lysed in tissue lysis buffer (100 mM Tris-HCl pH8.5, 5 mM EDTA, 200 mM NaCl, 0.2% SDS) + proteinase K 600 μg/ml (Roche) in a thermoshaker at 56°C and 400 rpm shaking overnight. Tissue lysis solution was centrifuged 5 min at 13,000 rpm and DNA in supernatant was precipitated by gently mixing with one volume of isopropanol. Precipitates were centrifuged 5 min at 13,000 rpm and supernatant was discarded. DNA pellets were washed once with two volumes of 75% ethanol. Pellets were air-dried and dissolved in nuclease-free water. Purified genomic DNA was used for genotyping PCRs for Cre transgene, caspase-8 flox allele, *Il-33* citrine allele and *St2* wt or KO alleles, using Mouse Kapa Genotyping kit (Merck-Millipore), primers (500 nM, Life Technologies) and nuclease-free water. PCR primers are listed in Supplementary Table 2. PCR products and 100 bp DNA marker (New England Biolabs) were loaded on a 2% agarose gel in Tris-Acetate-EDTA buffer, separated by electrophoresis and visualised using an imager (GeneSys Workstation).

### Cutaneous lesion scoring

Lesion scoring was calculated as the percentage and severity of Total Body Surface Affected (TBSA) by lesions. Abdominal Surface (AS) was considered as 30% of TBSA and Posterior Surface (PS), comprising back, flanks and head as 70% of TBSA. The Scoring system is presented in Supplementary Table 1. Both measurements were combined to obtain the final TBSA as follows:

TBSA %= 100 * (0.7*PS) +(0.3*AS)).

The experimental endpoint was reached for a TBSA of 50%.

### Survival measurement

Animals were monitored daily from birth for the development of cutaneous lesions as above. The experimental endpoint was reached for a TBSA of 50%. The survival rates were compiled and presented in a Kaplan-Meier curve. Statistical analysis is presented below.

### Mouse skin sections

Dorsal and abdominal skin biopsies were collected at indicated ages. For skin cryosections, samples were snap frozen and stored at -80 °C. Cryosections were cut in OCT at 7 μm thickness on Epredia CryoStar NX50 at -20°C chamber temperature and mounted onto lysine-coated Superfrost slides (VWR). Sections were fixed for 20 min in 4% ParaFormAldehyde (PFA) at RT prior to immunostaining. For paraffin embedded sections, tissue samples were fixed in 4% PFA overnight at 4 °C then stored at 4 °C in 100% Ethanol. They were then embedded in paraffin. Paraffin blocks were then sectioned at 7 μm thickness and mounted onto Lysine-coated slides and stained with Haematoxylin and Eosin (H&E) at the Histology Facility at University of Bristol.

### Epidermal thickness measurement

Pictures of skin sections stained with H&E as above were taken with 5X objective on Zeiss Apotome Axio Observer (Carl Zeiss) using the tile function of Zeiss Zen Blue software. Epidermal thickness was measured using the polygon function of ImageJ software (National Institutes of Health.

### Antibodies

The following primary antibodies were used: anti-P-MLKL (Abcam, 187091), anti-mIL-33 (R&D System, AF3626), anti-mTNF (BD Pharmingen, 559064), anti-cleaved caspase-3 (R&D Systems), anti-RIPK3 (Abcam,), anti-Krt14 (Neomarkers), anti-Krt10, Krt6 and loricrin (Biolegend), anti-CD3 (Agilent), anti-F4/80 (BD Biosciences, T45-2342) and anti-Gr1 (BD Biosciences, RB6-8c5). As secondary antibodies donkey-anti-mouse 488 AlexaFluor (1:2000), goat anti-rabbit 488, goat anti-rabbit 594, donkey anti-rat 594 and donkey anti-goat 594 were used in combination with DAPI as counterstaining for DNA. Fluorescent images were acquired on a Zeiss LSM800 confocal laser scanning microscope using ZEN 2.6 Software. DAPI was purchased from Sigma. Slides were mounted in fluorescence mounting solution Vectashield (H-1000, Vectastain Laboratories Inc.).

### Immunostainings

Double stainings for P-MLKL and IL-33 were performed on frozen skin section fixed in 4% PFA. Sections were blocked for one hour in PBS-0.05% Tween (PBS-T) + 10% FCS, then incubated overnight at 4°C with anti-P-MLKL antibody (1/1000) + anti-mIL-33 antibody (1/50) in PBS-T 10% FCS. Sections were washed 3x in PBS-T then incubated for one hour at room temperature with secondary antibodies Alexa488 anti Rabbit+ Alexa594 anti-goat in PBS-T 10% FCS, Sections were washed in PBS-T, incubated for 10 min in DAPI, washed twice with PBS-T then mounted using Vectashield fluorescence protective mounting solution mounting medium. The same protocol was used for CD3 (1/100), F4/80 (1/50), Gr-1 (1/20) and TNF (1/20) using PBS-T +10% Goat serum as blocking solution. Secondary antibodies were Alexa 594 anti-rat and Alexa 594 anti-Rabbit. For anti-cleaved caspase-3 and anti-RIPK3 immunostainings, sections were permeabilized using 1/100 trypsin solution in water for 30 min at 37 °C prior to immunostaining. Primary antibodies were diluted 1/1000 and 1/100 respectively in PBS-T 10% goat serum. then incubated the following day with secondary antibody Alexa 594 anti-Rabbit, stained with DAPI (Sigma), then mounted in Vectashield. Fluorescent images were acquired on a Zeiss LSM800 confocal laser scanning microscope using ZEN 2.6 Software.

### Epidermal markers immunostainings

For epidermal differentiation markers, formalin-fixed paraffin-embedded sections underwent deparaffinization and rehydration steps using xylene and successive baths of 90%, 75% and 50% ethanol. Heat-induced antigen retrieval using citrate buffer ph 6 was performed for 20 min. Sections were left to cool down then were blocked in 10% Goat serum in PBS-T for one hour. Sections were then labeled with primary antibodies diluted in PBS-T + 10% Goat serum against K14 (1/500) + K10 (1/100), Loricrin (1/100) or K6 (1/100) overnight at 4 °C. Sections were washed 3 times in PBS-then incubated for 1h at room temperature with secondary antibodies Alexa488 anti Rabbit for Loricrin and K6 and Alexa488 anti-Mouse +Alexa 594-anti Rabbit for K14/K10 double staining. Sections were washed with PBS-T, incubated for 10 min at RT with DAPI, washed twice with PBS-T then mounted using Vectashield (VectaLabs) mounting medium.

### Statistical analysis

All the data were analysed using GraphPad Prism 9.3.1. For the comparison of means between two different groups, a two-tailed student t-test was performed, and ANOVA was used for multiple groups analysis. The survival rate is presented as Kaplan-Meier survival curve and was compared by log-rank test. Data were presented as mean ± s.e.m and the differences were considered to be significant when p<0.05.

## Acknowledgments

We thank Dr. Stephen Hedrick (UCSD, U.S.A.) for kindly providing caspase-8 Flox mice. We thank Pr. Daniel Pinschewer (University of Basel, Switzerland) for kindly providing the St2-/-mice. We thank Pr. Andrew McKenzie (MRC, Cambridge, UK) and Dr. James MacLaren (Cardiff University, UK) for kindly providing the IL-33-/-citrine reporter mouse. We thank Dr. Timothy Hughes and Dr. Wiola Zelek for helpful discussions. We thank Michelle Somerville for helpful support with the imaging facility. We thank Pr. B. Paul Morgan for helpful discussions and continuous support.

## Author contributions

Conceptualization: MCB

Methodology: MCB, AFN, VS

Investigation: AFN, VS, FB

Visualization: AFN, VS, FB

Supervision: MCB

Funding: LM, AB, MB, EHC and MCB

Writing—original draft: VS and MCB

Writing—review & editing: AFN, EHC, LM, AB and MB

