## Supplementary material for "Epidermal IL-33 drives inflammation in necroptosis-induced skin inflammation by recruiting TNF-producing immune cells"

**This PDF file includes:**

Supplementary Text
Table S1
Figs. S1 to S4

**Supplementary Text**

Supplementary methods:

Low $Ca^{2+}$ mouse keratinocyte culture medium:

| | |
|---|---|
| KCl (Sigma) | 400 mg/L |
| $MgSO_4$-$7H_20$ (Sigma) | 200 mg/L |
| NaCl (Sigma) | 6.8 g/L |
| $NaHCO_3$ (Sigma) | 2.2 g/L |
| $NaH_2PO_4$-$H_2O$ (Sigma) | 140 mg/L |
| Glucose (Sigma) | 1 g/L |
| Non essential amino acids (Gibco) | 1X |
| Essential vitamins (Gibco) | 1X |
| Essential amino acids (Gibco) | 1X |
| Chelex-treated FCS | 4% |
| Antibiotics/antimycotics (Gibco) | 1X |
| mEGF (Peprotech) | 2.5 ng/ml |
| $CaCl_2$ (Sigma) | 45 μM |
| NaOH | 5 mM |
| Glutamine (Gibco) | 1% |

**Table S1. Lesion scoring method**

% Total Body Surface Affected (TBSA)

| | Severity | TBSA % |
|---|---|---|
| 0 | None | 0 |
| 1 | Faint | 1 – 10 |
| 2 | Low | 11 – 20 |
| 3 | Moderate | 21 – 35 |
| 4 | High | 36 – 45 |
| 5 | Very high (endpoint) | 46 + |

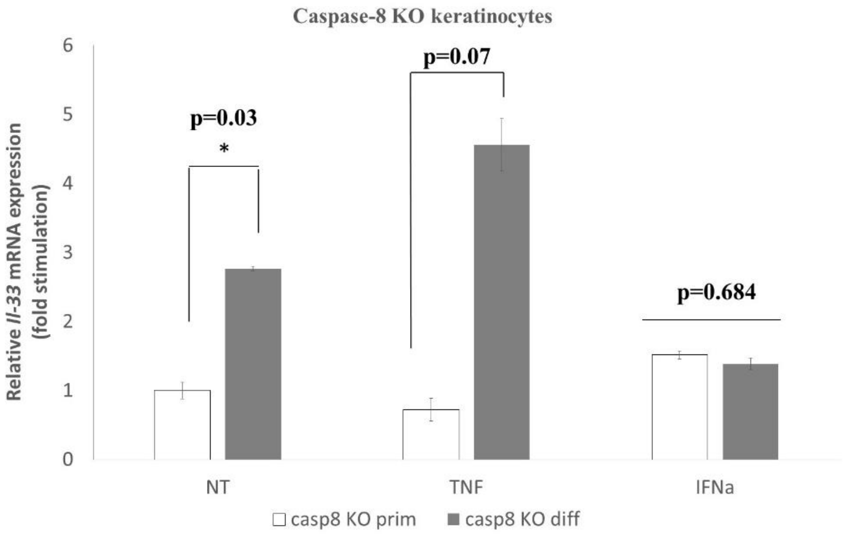


**Fig. S1.**

**Differentiation induces *Il-33* expression in caspase-8 KO keratinocytes.**
Representative data from three independent experiments.
Primary or differentiated caspase-8 KO keratinocytes were stimulated for 6h with mTNF (50 ng/ml) or mIFNα ($10^6$ IU/ml). RNA was isolated and Il-33 expression was quantified by qRT/PCR by comparison to *Gapdh* gene expression.

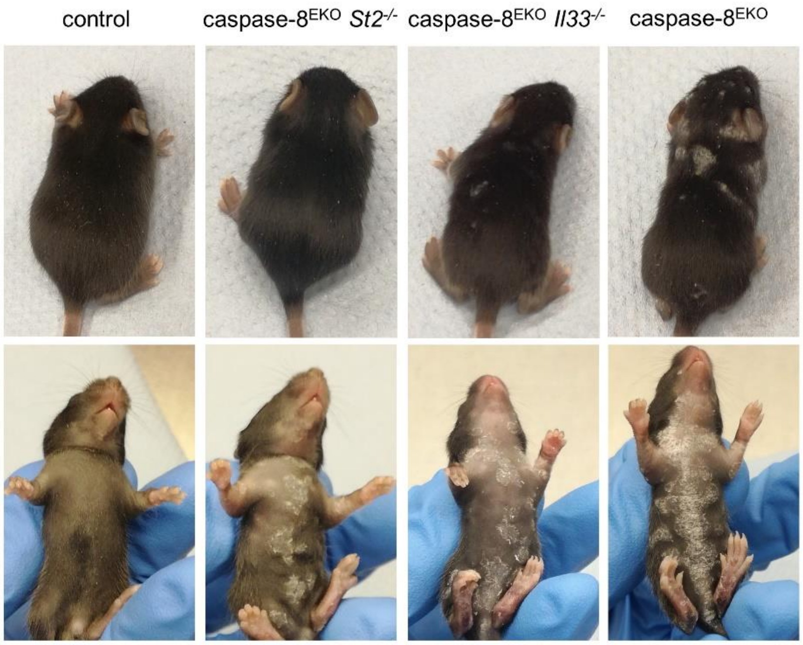


**Fig. S2.**
**Skin inflammation in caspase-8$^{EKO}$ *St2*$^{-/-}$ and caspase-8$^{EKO}$ *Il33*$^{-/-}$ mice at P12.**

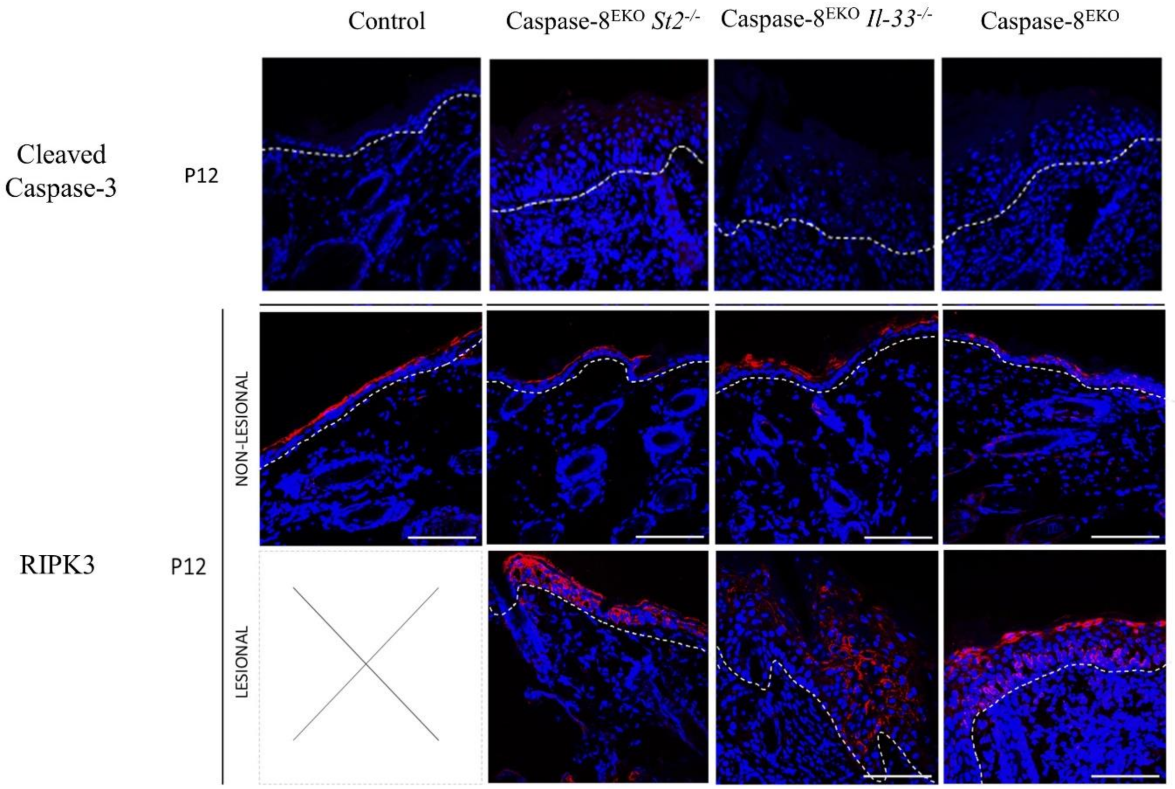


**Fig. S3.**

**Cell death markers in the skin of caspase-8$^{EKO}$ *St2*$^{-/-}$ and caspase-8$^{EKO}$ *Il33*$^{-/-}$mice.**

**A:** Cleaved Caspase-3 immunostaining in the skin of caspase-8$^{EKO}$ *St2*$^{-/-}$ and caspase-8$^{EKO}$ *Il33*$^{-/-}$ mice.

**B:** RIPK3 immunostaining in the skin of Caspase-8$^{EKO}$ *St2*$^{-/-}$and caspase-8$^{EKO}$ *Il33*$^{-/-}$ mice.

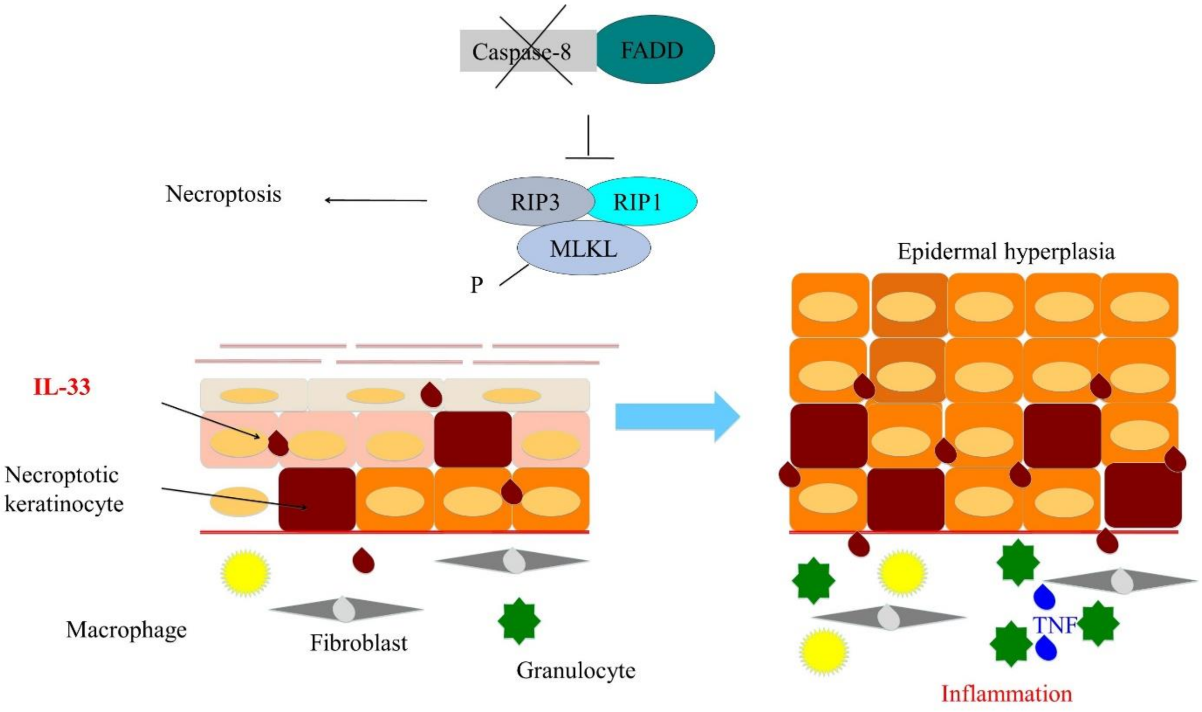


**Fig. S4**
**Model: IL-33 drives inflammation in necroptosis-induced skin inflammation.**
IL-33 released by necroptotic epidermal keratinocytes in the first days after birth recruit infiltrating granulocytes to the dermis which produce TNF, fuelling further keratinocyte necroptosis and cutaneous inflammation.
